# Prokaryotic winged helix domains as dsDNA adenine methylation sensors

**DOI:** 10.1101/2023.06.07.544091

**Authors:** Igor Helbrecht, Daniel Heiter, Weiwei Yang, Thomas Lutz, Laurence Ettwiller, Matthias Bochtler, Shuang-yong Xu

**Author notes:** These authors contributed to this work equally (Igor Helbrecht, Daniel Heiter). Corresponding authors Dr. Matthias Bochtler and Dr. Shuang-yong Xu.

## Abstract

Winged helix (wH) domains, also termed winged helix-turn-helix (wHTH) domains, are widespread in all kingdoms of life, and have diverse roles. In the context of DNA binding and DNA modification sensing, some eukaryotic wH domains are known as sensors of non-methylated CpG. In contrast, the prokaryotic wH domains in DpnI and phi.HhiV4I act as sensors of adenine methylation in 6mApT (6mA = N6mA) context. DNA binding modes and interactions with the probed dinucleotide are vastly different in the two cases. Here, we show that the role of the wH domain as a sensor of adenine methylation is widespread in prokaryotes. We present previously uncharacterized examples of PD-(D/E)XK—wH (FcyTI, Psp4BI), PUA—wH—HNH (HtuIII, Hsa13891I), wH—GIY-YIG (Ahi29725I, Apa233I) and PLD—wH (Aba4572I, CbaI) fusion endonucleases that sense adenine methylation in the Dam G6mATC, and possibly other, slightly more relaxed contexts. Representatives of the wH domain endonuclease fusion families with the exception of the PLD—wH family could be purified, and an *in vitro* preference for adenine methylation in the Dam context could be demonstrated. Like most other MDREs, the new fusion endonucleases except those in the PD-(D/E)XK—wH family cleave close to, but outside the recognition sequence. Taken together, our data illustrate the widespread combinatorial use of prokaryotic wH domains as adenine methylation sensors.

## Introduction

In most restriction-modification (R-M) scenarios, nucleobase modification serves as a mark of self and provides protection against endonuclease digestion. In some cases, however, phages have learnt to exploit this principle, by modifying their own DNA, either by incorporation of non-standard nucleoside triphosphates, or by post-replicative modifications catalyzed either by host or phage enzymes. Modification dependent restriction endonucleases (MDREs) specifically target such modified DNA (modified base or backbone). The MDREs come in two main groups, distinguished by the presence or absence of nucleoside triphosphate (NTP) consuming motor proteins. The NTP independent proteins are typically modular, with separate modification sensing and DNA cleavage domains. Because of this architecture, DNA cleavage typically takes place at a distance from the site of modification. For some enzymes a single site is sufficient, but typically, cleavage is most efficient when it is directed by appropriately spaced modifications, which cooperate to position an endonuclease dimer for a double strand (ds) cut in the DNA.

The catalytic domains present in restriction can be grouped into the almost universally used hydrolases and the very rarely used lyases (1). The hydrolases in turn can be grouped into a surprisingly small set of phylogenetically unrelated enzyme groups. PD-(D/E)XK enzymes are named for characteristic amino acids (aa) built around a central β-sheet, which harbors one or two catalytic Mg^2+^ ions (2). The metal ions are held in place in part by the D and (D/E) moniker residues, which together with the moniker K residue activate a water molecule for direct inline attack on the scissile phosphate (3,4). HNH enzymes, also called ββα-Me enzymes or His-Me finger enzymes (5,6), harbor a single metal cation in their active site. Metal identity requirements are less strict than for PD-(D/E)XK enzymes. Many divalent transition metal ions are acceptable (7). Like PD-(D/E)XK enzymes, the HNH enzymes are believed to catalyze attack on the scissile phosphate by a water molecule. However, water activation is not by a lysine residue, but by the first histidine of the HNH motif (8,9). GIY-YIG enzymes also bind a single metal cation in the active site. These enzymes activate the water molecule by a tyrosine residue, most likely from the GIY motif (10). Finally, there are also completely metal-independent endonuclease domains. They resemble phospholipase D, therefore the enzymes containing them are known as PLD endonucleases (11). The PLD enzymes are believed to catalyze phosphodiester cleavage via a covalent intermediate (12).

The modification sensor domains, like the endonuclease domains, are now understood to be classifiable into only a few groups of phylogenetically unrelated sensors. The largest group of sensors is the PUA superfamily (13). PUA superfamily sensors comprise SRA domains with specificity for 5mC, as in MspJI (14), and related domains (15,16), originally also termed SRA domains, with specificity for 5-hydroxymethylcytosine (5hmC) and glucosyl-5-hydroxymethyl-cytosine (g5hmC), as in the PvuRts1I family of restriction endonucleases (17,18). The PUA superfamily also comprises EVE domains specific for 5mC and 5hmC, as found in VcaM4I (19), and YTH domains (Yth-McrB fusion) specific for 6-methyladenine (6mA) (20-22), as well as ASCH domains. Bioinformatic analysis has suggested that ASCH domains might be 6mA readers (23), but this prediction is not confirmed by experimental data so far. Instead, it has been shown that the *E. coli* protein YqfB, an ASCH domain protein, is able to hydrolyse various N4-acylated cytosines (4acC) and cytidines (24). All PUA superfamily domains are engaged in nucleotide flipping. Irrespective of their detailed specificity, they scrutinize the modified base in a dedicated pocket of the enzyme (25).

Apart from the PUA superfamily, other modification sensor domains may also be involved in restriction, such as the NEco domain in EcoKMcrA with affinity for 5mC and 5hmC (26). Unlike the PUA superfamily domains, the NEco domain senses 5mC or 5hmC without nucleotide flipping, in the context of dsDNA (27). Finally, a winged helix (wH) domain has been described as a 6mA sensor in DpnI. Like the NEco domain, the wH domain senses nucleobase modifications in the context of dsDNA, without flipping (28). However, in contrast to the NEco domain, it has specificity for 6mA rather than 5mC. Also, in contrast to NEco, which recognizes methyl groups of fully methylated CpG in two separate pockets, the wH domain recognizes methyl groups of fully methylated ApT in a single pocket, exploiting their proximity in space. The wH domain in DpnI is unusual in being fused to a nuclease domain which has separate sequence (GATC) and modification (6mA) specificity (29). Therefore, it acts more like an effector domain in Type IIE enzymes, except that both the nuclease and sensor/effector domain are specific for methylated rather than non-methylated DNA.

Winged helix (wH) domains are a group of DNA binding domains that belong to the superfamily of helix-turn-helix (HTH) proteins (30-32). Structurally, canonical winged helix domains consist of an N-terminal α-helix and β-strand, the HTH-motif, and a β-hairpin. The “wings” of the motif are the loops connecting the strands of the β-hairpin and immediately downstream of it (28). Winged helix motifs were first found in transcription factors, but it is now clear that they also have roles in transcription initiation complexes (33), in the binding of left-handed Z-DNA (34) or RNA (35), or in protein-protein interactions (36). In transcription factors, wH domains tend to interact with DNA just as would be expected for the HTH motif that is embedded within them. In other words, they insert the second helix of the HTH motif, which is the third helix of the wH domain, into the major groove of DNA (31). However, other DNA binding modes are also possible in special cases (37,38). A recent example for such alternative binding modes is the complexes of eukaryotic winged helix domains with dsDNA containing non-methylated CpG (39-41). A winged helix motif in a restriction endonuclease (REase) was first noticed in the DNA binding domain of FokI (36), but this particular wH domain does not appear to be involved in DNA interactions.

A role of the winged helix domain in adenine methylation sensing was first noticed in DpnI. DpnI is a G6mATC specific endonuclease, that cleaves within the recognition sequence and has a strong preference for DNA that is adenine methylated in both strands (29). In DpnI, the winged helix domain plays the role of an effector domain that senses 6mA separately from the PD-(D/E)XK nuclease domain (with slightly relaxed sequence specificity compared to the nuclease domain) (28) **(Fig. 1B, C)**. More recently, a winged helix domain was also implicated in the sensing of adenine methylation, also in the G6mATC context, in HHPV4I (phi.HhiV4I (42). Phi.HhiV4I (a.k.a. HHPV4I) is a three-domain enzyme, with a PUA (SRA)-like domain at the N-terminus, a winged-helix domain in the middle, and an HNH endonuclease domain at the C-terminus. The PUA superfamily domain, described as an SRA domain by Lu and colleagues, appears not to be involved in DNA modification sensing (42). By contrast, the winged helix domain cleaves Dam^+^ preferentially over Dam^-^ DNA, and it has much higher affinity to Dam^+^ than to Dam^-^ DNA in gel shift experiments. Unlike DpnI, phi.HhiV4I cleaves at a distance from the site of adenine methylation, suggesting that the endonuclease domain is directed by the winged helix domain and does not sense adenine methylation on its own (42).

**Fig. 1.**
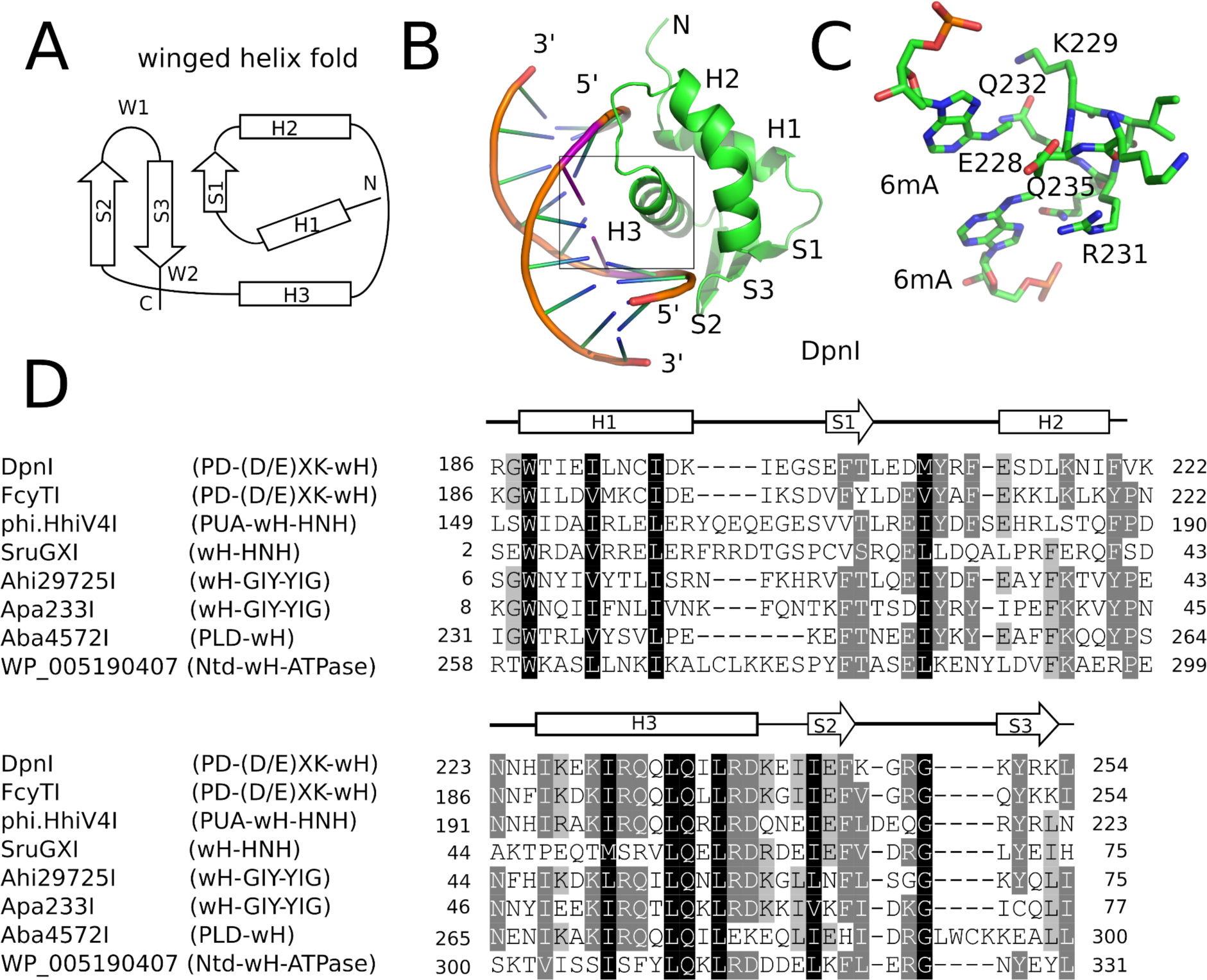
Winged helix (wH) domain fold and role as a methylation sensor. **A.** Canonical wH fold. **B.** Cartoon representation of the DpnI wH domain bound to fully methylated DNA, based on the crystal structure (28). **C.** Methyl binding region of the DpnI wH domain. **D.** Alignment of representative winged helix domains in endonuclease or NTPase fusions. The secondary structure annotation is based on the DpnI experimental structure, analyzed for secondary structure elements using dssp (dsspweb.html). Arrows indicate residues involved in 6mA recognition.

In this work, we show that the winged helix domain is used widely as an adenine methylation sensor in MDREs **(Fig. 1D)**. We present additional examples of proteins that share the PD-(D/E)XK—wH architecture with DpnI, or the PUA—wH—HNH architecture with HHPV4I. Additionally, we show that the wH domain is also naturally paired with an HNH domain in the absence of a PUA-superfamily domain, with a GIY-YIG endonuclease domain, with a PLD (phospholipase D) domain, or with an NTPase (GTPase/ATPase) domain. For the PD-(D/E)XK—wH, PUA—wH—HNH, wH—HNH, wH—GIY-YIG, and PLD—wH enzymes, we detect Dam^+^ dependent toxicity in *E. coli* cells, suggesting that these enzymes have adenine methylation dependent MDRE activity. For the PD-(D/E)XK—wH, PUA—wH—HNH, wH— HNH, and wH—GIY-YIG, but not the PLD—wH enzymes, we find representatives that are active also in *in vitro*, and we show that their preferred substrate is fully methylated DNA with one enzyme exception. Unlike DpnI, many of the new MDREs cleave DNA outside the G6mATC recognition sequence.

## Materials and Methods

### Materials

*E. coli* T7 expression strains C2566 (Dam^+^) and its isogenic Dam-deficient strain ER2948 (constructed and provided by Dr. Lise Raleigh, New England Biolabs), expression vector pTXB1, pBR322, phage λ DNA (Dam^+^ or Dam^-^), 2-log (1 kb plus) DNA ladder, chitin beads, restriction enzymes, EcoGII methylase (frequent adenine methylase), Q5 DNA polymerase PCR master mix and cloning kit (Hi-Fi DNA assembly enzyme mix), NEBExpress Ni-NTA magnetic beads, and dZTP (2-aminoadenine triphosphate is abbreviated as base Z in the literature, in this paper we use dZ to differentiate from left-handed Z DNA) were provided by New England Biolabs (NEB). Ni-agarose beads were from Qiagen or NEB. The T7 expression vector pET21b with C-terminal 6xHis tag was originally purchased from Novagen (NdeI-XhoI). 5hmdCTP/dGTP/dATP/dTTP mix was purchased from Zymo Research. FPLC DEAE, Heparin columns (5 ml) were purchased from GE HealthCare or Cytiva.

### Digests

Restriction digests were carried out in 1x NEB restriction buffers: buffer 1.1 (10 mM Bis-Tris-Propane-HCl, 10 mM MgCl_2_, 100 µg/ml BSA or recombinant albumin, pH 7.0 at 25 °C); 2.1 (50 mM NaCl, 10 mM Tris-HCl, 10 mM MgCl_2,_ 100 µg/ml BSA or recombinant albumin, pH 7.9 at 25 °C); 3.1 (100 mM NaCl, 10 mM Tris-HCl, 10 mM MgCl_2,_ 100 µg/ml BSA or recombinant albumin, pH 7.9 at 25 °C) or CutSmart buffer (50 mM Potassium Acetate, 20 mM Tris-acetate, 10 mM Magnesium Acetate, 100 µg/ml BSA or recombinant albumin, pH 7.9 at 25 °C). For restriction digestions in different divalent cations, a medium salt buffer (50 mM NaCl, 20 mM Tris-HCl, pH 7.5) was supplemented with divalent cations in 0.1, 1, and 10 mM final concentration.

### Synthetic oligos with modified and unmodified GATC sites

(Single-stranded DNA oligos were synthesized by NEB Organic synthesis division): Top strand Gm6ATC (top M+)

5’/56FAM/ACTCATGCAGGCATGCAGG/m6A/TCGCAGTCAGATTTATGTGTCATATAGTACGTGATTCAAG 3’

Bottom strand Gm6ATC (Bottom M+)

5’CTTGAATCACGTACTATATGACACATAAATCTGACTGCG/m6A/TC CTGCATGCCTGCATGAGT 3’

Top strand GATC (unmodified. Top M-) 5’/56FAM/ACTCATGCAGGCATGCAG<u>GATC</u>GCAGTCAGATTTATGTGTCATATAGTACG TGATTCAAG 3’

Bottom strand GATC (unmodified. Bottom M-) 5’CTTGAATCACGTACTATATGACACATAAATCTGACTGC<u>GATC</u>CTGCATGCCTGCATG AGT 3’

Duplex oligos abbreviation:

Fully modified = M+ top/M+ bottom = M+/M+

Unmodified = M-top/M-bottom = M-/M-

Hemi-modified = M+ top/M-bottom = M+/M-

Hemi-modified = M-top/M+ bottom = M-/M+

Duplex oligos were digested by DpnI (2 U), MboI (5 U), FcyTI (0.1 µg), Ahi29725I and Apa233I (1 µg) in NEB buffer 2.1 at 37°C for 1 h. For phi.HhiV4I digests reactions were carried out in NEB buffer 2.1 supplemented with 1 mM MnCl_2_ (this enzyme is a Mn^2+^ -dependent REase, see below).

Duplex oligos and protein concentration in restriction digestions:

The duplex oligos concentration is approximately 18 nM (60mer, 21 ng in 30 µl total volume). The protein concentration is calculated below:

Ahi29725I protein (dimer) MW = 24.62 x2 = 49.24 kDa.

Ahi 0.1 µg = 68 nM

Ahi 1 µg = 677 nM

Apa233I protein (dimer) MW = 24.17 x2 = 48.34 kDa

Apa 0.1 µg = 69 nM

Apa 1 µg = 690 nM

phi.HhiV4I protein (dimer) MW = 44.67 x2 = 89.34 kDa

phi.HhiV4I 0.1 µg = 37 nM

phi.HhiV4I 1 µg = 373 nM

FcyTI protein (dimer) MW = 30.76 kDa x2 = 61.52 kDa

FcyTI 0.1 ug = 54 nM

FcyTI 1 ug = 542 nM

### Protein expression

C2566 competent cells (cloning grade) were provided by NEB. ER2948 competent cells were prepared by a modified rubidium chloride method. *E. coli* cells were usually cultured at 37 °C to mid log phase, and IPTG was added to the culture at 0.5 mM final concentration for protein production (at 18 °C overnight). Cells were lysed by sonication in chitin column buffer or Ni-agarose column buffer. Clarified cell lysates with over-expressed proteins (target protein-intein-CBD) or C-terminal 6xHis tagged protein were loaded onto chitin or Ni-agarose columns respectively for affinity purification. The protein purification protocols were used as recommended by the manufacturers. In some cases, the partially purified proteins were further purified by chromatography through DEAE (flow through to remove nucleic acids at 0.3 M NaCl concentration) and Hi-Trap Heparin (GE HealthCare or Cytiva, 5 ml). The plasmid preparation and transformation protocols followed NEB recommendations. BigDye® Terminator v3.1 Cycle Sequencing Kit was purchased from Thermo-Fisher (Applied Biosystems). Restriction gene inserts in plasmids were sequenced to verify correct sequences. All synthetic genes (gene blocks) are listed in the Suppl. Materials. Dam+ pBR322 DNA fragments after restriction digestion were sequenced to determine the cut sites. DNA sequence edits were carried out using DNAStar or Geneious software packages. BlastP searches in GenBank and UniProt databases were performed using the respective web servers. NCBI Pfam and conserved domains were used to visualize protein domains of REase homologs.

### CLANS analysis

For CLANS analysis, the homologs of the five groups of wH-containing enzymes (sequences of the reference proteins are listed in the Suppl. Materials) were obtained by blast (BlastP, default settings in the UniProtKB website) using the UniProtKB reference proteomes and Swiss-Prot database. The resulting homolog protein fasta files were combined and subjected to CLANS analysis using the MPI Bioinformatics Toolkit. The result of the CLANS analysis is visualized with the CLANS java application.

### Phylogenetic analysis

To perform phylogenetic analysis, we first built a hmm profile with the wH domains of the six representative wH-containing enzymes (sequences of the reference proteins are listed in the Suppl. Materials). The resulting profile was used to search homologs using hmmsearch (HMMER v3.1b2) on the combined fasta file (see CLANS analysis). We extracted the wH domain in each homolog protein and performed multiple alignment together with the five representative wH domain using MAFFT (v7.508). The maximum likelihood tree was constructed using RAxML (v8.2.12) and visualized with iTOL (https://itol.embl.de/).

## Results

### Bioinformatic screen of wH domain endonucleases

Some wH domains are adenine methylation (6mA) specific. This notion was originally suggested by the demonstration of 6mA specificity of the DpnI wH domain and further strengthened by the observation of adenine specificity of the wH domain of HHPV4I, which was reported when this work was being finalized. In the hope to find new adenine methylation specific endonucleases, we searched for fusions of wH domains with nuclease domains known to play roles in restriction modification. Apart from additional PD-(D/E)XK—wH and PUA— wH—HNH endonucleases, we also identified proteins with a wH—HNH, wH—GIY-YIG, and PLD—wH architecture. In addition, we found PD-(D/E)XK—wH—NTPase cases, that can be considered as fusions of a DpnI-like protein with an McrB-like NTPase (GTPase/ATPase) domain, and NTD—wH—NTPase cases, apparently without a nuclease domain, and with an N-terminal domain of unknown function (sequences not shown). On additional example is the wH—Mrr catalytic (PD-QxK)—NTPase fusion endonuclease (sequence not shown). For our studies, we concentrated on the NTP-independent enzymes with wH and endonuclease domains. In CLANS analysis of the full-length proteins (with BamHI (GGATCC) and related enzymes as a control), the wH-domain containing endonucleases segregated clearly into separate groups, driven by the sequence similarity within the endonuclease groups **(Fig. 2)**.

**Fig. 2.**
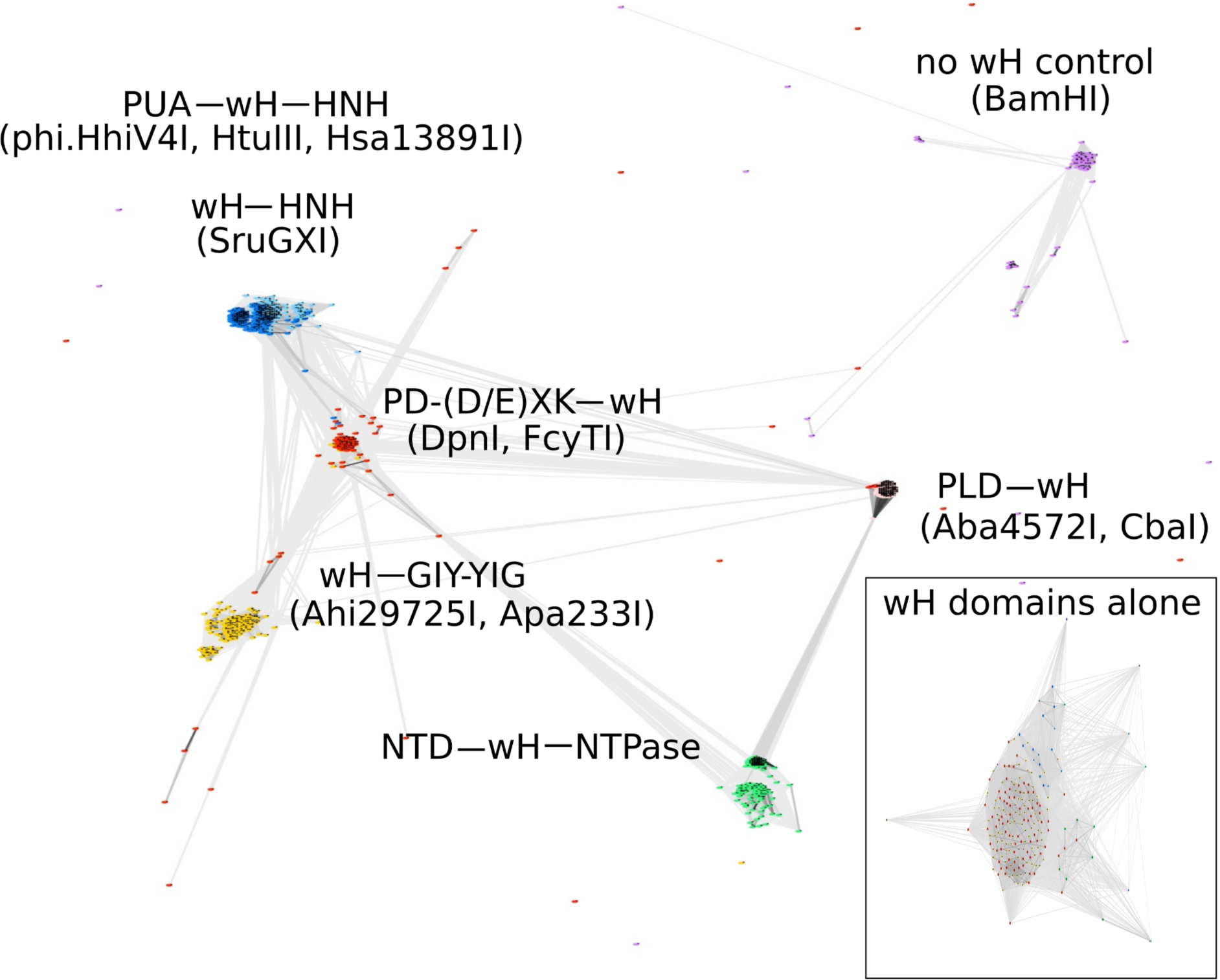
CLANS analysis of the full-length wH fusion proteins with PD-(D/E)XK (marked as red), PUA and HNH (dark blue); HNH (light blue), GIY-YIG endonuclease domains (yellow) and NTD (N-terminal domain) and NTPase (green) studied in this work, with BamHI and similar enzymes as a control group (purple). BOX: CLANS analysis of the wH domains alone, color-coded as in full-length proteins.

However, when we limited the CLANS analysis to the wH helix domains alone, the segregation into groups according to the endonuclease domain was not so clear. In particular, wH domains from PD-(D/E)XK and GIY-YIG endonucleases were fully intermingled, possibly suggesting multiple separate fusion events **(Fig. 2 box)**.

### Bioinformatic evidence for MDRE activity

Adenine methylation in bacteria occurs frequently in GATC context, the target sequence of the Dam methyltransferase (MTase) (43), which is widely distributed in bacteria because of its diverse house-keeping roles, including in bacterial DNA replication (44) and mismatch repair (45,46). Hence, it was likely that the putative MDREs with wH domain might detect adenine methylation in this sequence context. This idea was further supported by the precedent of the wH domain in DpnI, which is known to be adenine methylation specific, provided the adenine methylation is present in the Dam context (with some leeway for the outer bases). Inspection of the crystal structure of the winged helix domain of DpnI in complex with target DNA revealed that the same residues contribute to both the methyl binding pocket and the sequence specificity, suggesting that methyl detection and detection of the G6mATC target sequence are intricately linked. With the exception of SruGXI, the representative of the wH—HNH endonucleases, the residues that are involved in methylation and sequence sensing are conserved in the wH domains of the new putative MDREs **(Fig. 1D)**. It was therefore likely that the new candidate MDREs would also display specificity for G6mATC.

If the new candidate MDREs were specific for adenine methylation in the G6mATC context, they should not co-occur with Dam-like adenine MTases to avoid self-restriction. We tested this prediction for 899 wH endonuclease domain fusions, by inspecting the genomic neighborhood within a 10 kb interval. PD-(D/E)XK—wH domains co-occurred with predicted C5 methyltransferases in 40 cases. In some cases, they also co-occurred with a predicted 4mC (N4mC) or 6mA MTase directly adjacent to it. In these cases (e.g. *Bacteroidota* bacterium isolate, CP064983.1, *Moraxella ovis* strain CP011158.1), the putative DNA MTase was inactivated by a frame shift. The PUA—wH—HNH and wH—HNH co-occurred in 47 cases with Eco57I-like MTases (of Type IIG R-M-S fusion enzymes). These MTases are predicted to be 6mA MTases, but since their expected target sequence is CTGA<u>A</u>G (site of methylation underlined), no conflict with a G6mATC specific MDRE is expected. In the case of the wH— GIY-YIG endonucleases, we found four instances of a EcoRI-like adenine MTase nearby. As these MTases are expected to methylate GA<u>A</u>TTC, again there is no conflict. Finally, for the PLD—wH endonucleases, we detected 17 cases of proximity to EcoEI-like (GAGN7ATGC) or EcoR124I-like (GA<u>A</u>N6RTCG) Type I methyltransferases, also causing no conflict. Taken together, the data show that most of the putative MDREs with wH domain are stand-alone enzymes, which are not associated with Dam-like methyltransferases. Remarkably, this was not the case for stand-alone winged helix domains, which could have Dam-like MTases in their neighborhood. In these cases, toxicity would not be expected because of the absence of a nuclease domain. There are a few wH fusion endonucleases in the wH-GIY-YIG family and PLD-wH family that co-exist with Dam-like methylases. How the hosts avoid self-restriction is unknown. Taken together, the bioinformatic data are consistent with the hypothesis that the wH MDREs are specific for Dam methylated DNA.

### The wH fusion proteins exhibit Dam^+^ dependent toxicity to *E. coli* cells

If the putative MDREs were specific for Dam methylated DNA, they should be more toxic to Dam^+^ (C2566) than to Dam^-^ (ER2948) *E. coli* cells. We tested this prediction with our IPTG inducible expression constructs, both under basal (0 mM IPTG) conditions, with only minimal leaking protein expression, and under induction conditions (0.5 mM IPTG). As Dam^-^ cells have a considerable fitness disadvantage compared to Dam^+^ cells, and since Dam^-^ competent cells were roughly an order of magnitude less competent than Dam^+^ cells, we avoided direct comparisons between Dam^+^ and Dam^-^ cells. Instead, we quantified the reduction in colony counts for transformations with expression plasmid compared to colony counts for transformation with empty vector. For Dam^-^ ER2948 cells, the colony count ratio was about 1:1, within experimental error. For Dam^+^ C2566 cells, the ratio was also not significantly different from 1:1, except in the case of CbaI, which caused a reduction in colony count by about two orders of magnitude even in the absence of IPTG induction. With the exception of the Aba4572I expression construct, the expression plasmids for all other putative MDREs caused a reduction in colony count by two to three orders of magnitude compared to the empty vector control in Dam^+^ C2566 cells under induction conditions. The experiment suggests that the wH endonuclease fusion proteins cleave Dam^+^ but not Dam^-^ DNA, or bind tightly to the modified sites and prevent DNA replication.

### PD-(D/E)XK—wH endonucleases

As representatives of the PD-(D/E)XK—wH family, we selected Psp4BI and FcyTI (GenBank accession numbers WP_102090895 and WP_094411979). The two enzymes have 58.4% and 58.7% amino acid (aa) sequence identity to DpnI, respectively. Psp4BI was chosen for *in vitro* characterization, because the source organism is psychrophilic, suggesting that the enzyme might be susceptible to heat inactivation at low temperatures, which would be desirable for biotechnological applications. The synthetic genes with *E. coli* optimized codons were cloned into pTXB1 in fusion with intein and CBD (chitin binding domain) and expressed in the Dam-deficient T7 expression strain ER2948. The two enzymes were affinity purified on a chitin column and released from the column by DTT triggered cleavage. The yield of Psp4BI was low due to poor expression of the fusion protein (Psp4BI-intein-CBD); Partially purified Psp4BI gave rise to a partial digestion pattern which was retained after 4 h at 25 °C - 37 °C (data not shown). By contrast, purified FcyTI was active on Dam^+^ pBR322 **(Fig. S1)**. FcyTI specific activity was determined as approximately 32,000 U/mg protein in buffer 2.1. High concentration of the enzyme inhibited restriction activity, probably as a result of protein aggregation (data not shown). FcyTI could be inactivated by heating at 65 °C for 30 min, which is a useful enzyme property for recombinant DNA applications **(Fig. S2)**. FcyTI endonuclease was originally found in the genome of *Flavobacterium cyanobacteriorum* that can grow at 20°C to 30°C.

Due to its better biochemical properties, FcyTI was used for *in vivo* toxicity study and modified oligos digestion. Run-off sequencing demonstrated that the enzyme was able to cleave within the G6mATC recognition sequence **(Suppl. Fig. S3)**.

To compare the activity of the enzyme towards fully methylated, hemi-methylated, and non-methylated DNA, we digested synthetic DNA oligoduplexes and quantified substrate and product amounts after restriction digestion by capillary electrophoresis (CE). The results showed that FcyTI was most active on fully methylated DNA, but also had partial activity on hemi-methylated DNA, similar to DpnI. No digestion product was detected for non-methylated DNA. MboI used as a control digested only unmodified GATC oligos. Fully and hemi-modified substrates were resistant to its activity (**Fig. 4**).

**Fig. 3.**
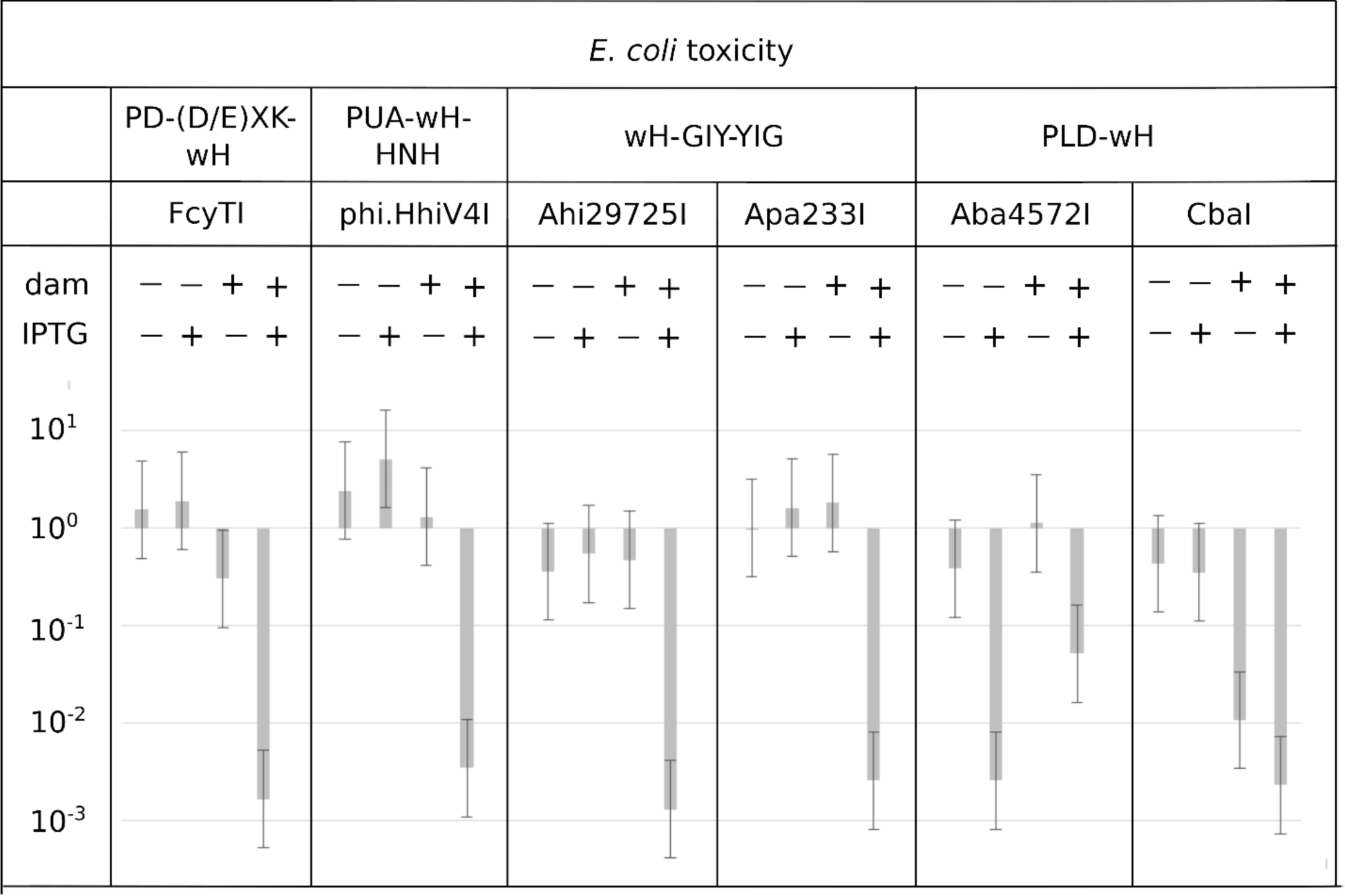
Toxicity of selected wH domain containing endonucleases to a Dam^+^, but not a Dam^-^ host. Expression vectors containing open reading frames for putative MDREs or empty plasmid (50 ng) were transformed into Dam positive C2566 (+) or Dam negative ER2948 (-) *E. coli* cells, with (+) or without (-) IPTG induction. The reduction in colony count for expression plasmid compared to empty vector, plotted on the ordinate, is a measure of toxicity.

**Fig. 4.**
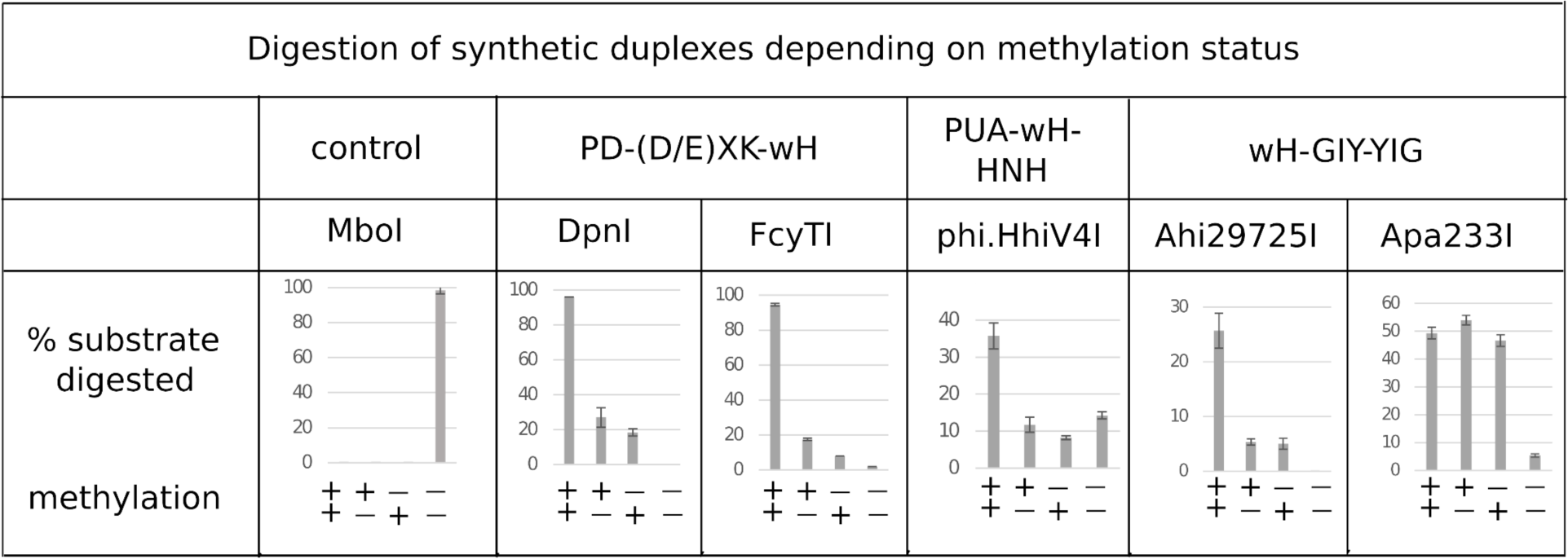
Digestion of a synthetic DNA oligoduplex depending on top and bottom strand methylation status in the G6mATC sequence context. 5 U of MboI (GATC) and 2 U of DpnI (G6mATC) were used as controls. FcyTI input was 0.1 μg protein (∼3 U) (higher concentration could obscure the digestion of hemi-modified duplex oligos). For three other enzymes, the amount of protein was 1 μg.

### PUA—wH—HNH and wH—HNH endonucleases

Consistent with the findings of Lu and colleagues (42), we observed that purified phi.HhiV4I was much more active in the presence of Mn^2+^ ions than in the presence of other divalent metal cations **(Suppl. Fig. S4).** Phi.HhiV4I showed low DNA nicking activity in Mg^2+^ buffer.

In agreement with the toxicity experiments **(Fig. 3)** and the results of Lu and colleagues (42), we found that the enzyme had higher activity against Dam^+^ than Dam^-^ pBR322, pUC19, λ DNA, and synthetic duplex oligos. If phi.HhiV4I cleaved at or near Dam^+^ sites, its cleavage products should be of similar size as those of DpnI digestion, and discrete bands (as opposed to a smear on the gel) should be observed. In our experiments, we saw only a partial match of fragment sizes, likely due to incomplete digestion **(Fig. 5A)** (see below for two-sites requirement for efficient cleavage). Dam^+^ phage λ DNA was also only partially digested while Dam^-^ λ DNA was not cut at all (λ DNA was partially methylated by the Dam methylase during rapid phage replication in *E. coli*) (**Fig. 5B**). When Dam^-^ λ DNA was methylated *in vitro* by Dam methylase or M.EcoGII, the DNA substrates now became cleavable by phi.HhiV4I (**Fig. 5C**), further demonstrating that GATC methylation is required for restriction. M.EcoGII-modified λ DNA appeared to be a slightly better substrate for phi.HhiV4I restriction, probably as a result of small enhancement of frequently modified adenine.

**Fig. 5.**
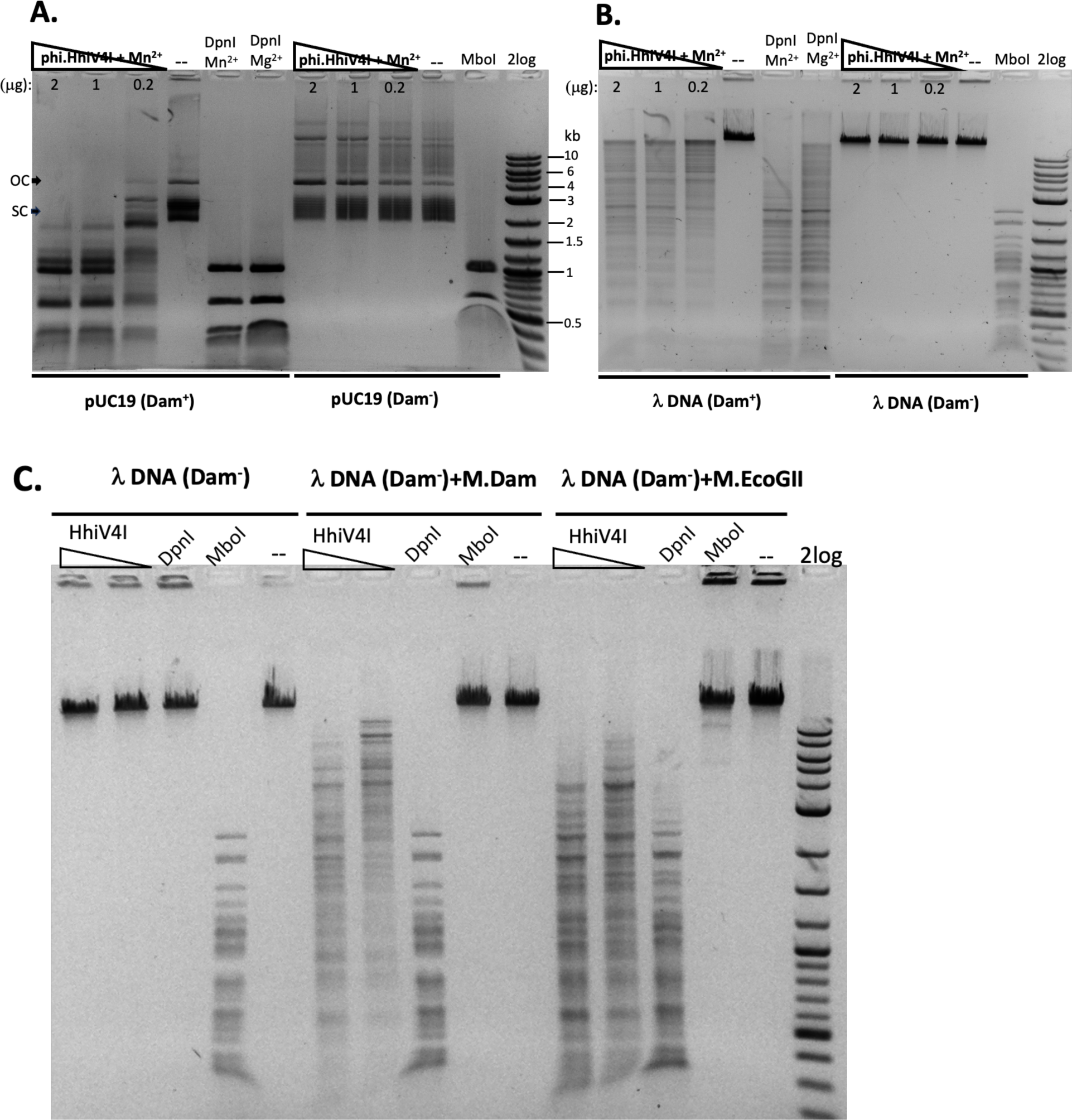
Methylation dependence of phi.HhiV4I. **A.** Restriction digests of Dam^+^ or Dam^-^ pUC19 in Mn^2+^ buffer. **B.** Restriction digests of Dam^+^ or Dam^-^ phage λ DNA in Mn^2+^ buffer (λ DNA partially modified by the host Dam methylase). DpnI (2 U) cleaves Dam^+^ DNA only. MboI (5 U) cleaves unmodified DNA only. **C**. phi.HhiV4I (HhiV4I) digestion (2 and 0.2 μg) of *in vitro* modified λ DNA (Dam^-^) by Dam methylase or frequent adenine methylase M.EcoGII in Mn^2+^ buffer. Following methylation reactions, the methylases were inactivated by heating at 65°C for 20 min. The DNA was then diluted for restriction digestion in Mn^2+^ buffer.

We could digest non-methylated DNA with excess phi.HhiV4I, suggesting that the dependence of the enzyme on adenine methylation was not absolute. This conclusion was confirmed with digestion of synthetic DNA with defined adenine methylation status. As expected, phi.HhiV4I was most active on fully methylated DNA, but had some activity on hemi- and non-methylated DNA **(Fig. 4)**. In agreement with the findings by Lu and colleagues (42) we saw no activity of phi.HhiV4I towards PCR products, which contained 5mC or 5hmC instead of C, in conditions conducive to digestion of 6mA containing DNA **(Suppl. Fig. S5)**. Since the PCR products contain 5mC and 5hmC in many different contexts, this result suggests that the enzyme has no activity against methylated or hydroxymethylated DNA, despite the presence of the PUA (SRA-like) domain. Phi.HhiV4I shows no activity on dZ (2-aminoadenine)-modified PCR either (**Suppl. Fig. S5**). Possible activity against WT T4 (g5hmC), phage 9g (dG+, deoxyarchaeosine), phiW-14 (pu-dT, putrescinylthymine) modified DNAs remains to be tested.

Phi.HhiV4I prefers to cut between two G6mATC sites with optimal spacers of 13-27 bp in Dam^+^ pBR322. Shorter spacers of 8-11 bp or longer spacers >42 bp were cleaved more slowly. Run-off sequencing of Dam^+^ phi.HhiV4I DNA confirmed that the enzyme cleaved in the vicinity of, but not within the G6mATC sequence, as previously reported **(Suppl. Fig. S6)**.

In contrast to the prophage encoded phi.HhiV4I, most PUA—wH—HNH enzymes, wH— HNH endonucleases are bacterial/archaea enzymes. For 15 of these enzymes and phi.HhiV4I as a positive control, we attempted expression in the Dam^-^ *E. coli* cells. Moreover, we quantified the ratio of colonies after transformation into Dam^+^ (C2566) and Dam^-^ (ER2948) cells. Restriction activity was examined in the presence of IPTG induction to elevate the genome conflict. Some restriction genes such as phi.HhiV4I and SruGXI had a strong toxic effect, as detected by 100 to 1000-fold reduction in transformation efficiency. Other ORFs caused an approximately 10-fold reduction in transformation efficiency consistent with partial restriction (+/-). The transformation of the HhaN23I gene caused the formation of very small colonies in the presence or absence of IPTG indicating partial restriction. A few ORF constructs showed no difference in transformation efficiency into Dam^+^ hosts, presumably as a result of poor expression or null activity (e.g. HboP9I). As a control, the pTXB1 empty vector could be readily transferred into C2566 (Dam^+^) or ER2948 (Dam-) cells in the presence of IPTG. The transformation efficiency of the empty vector into Dam^-^ cells was 5 to 10-fold lower than Dam^+^ cells probably due to two different methods of chemical treatments (C2566 cells are commercial grade competent cells from NEB and ER2948 cells were prepared in the lab by RbCl_2_/CaCl_2_ treatment) or due to the poor growth fitness of the Dam^-^ host (Dam modifications are involved in the initiation of *E. coli* DNA replication) **(Table 1 and Suppl. Fig. S7)**.

**Table 1:**
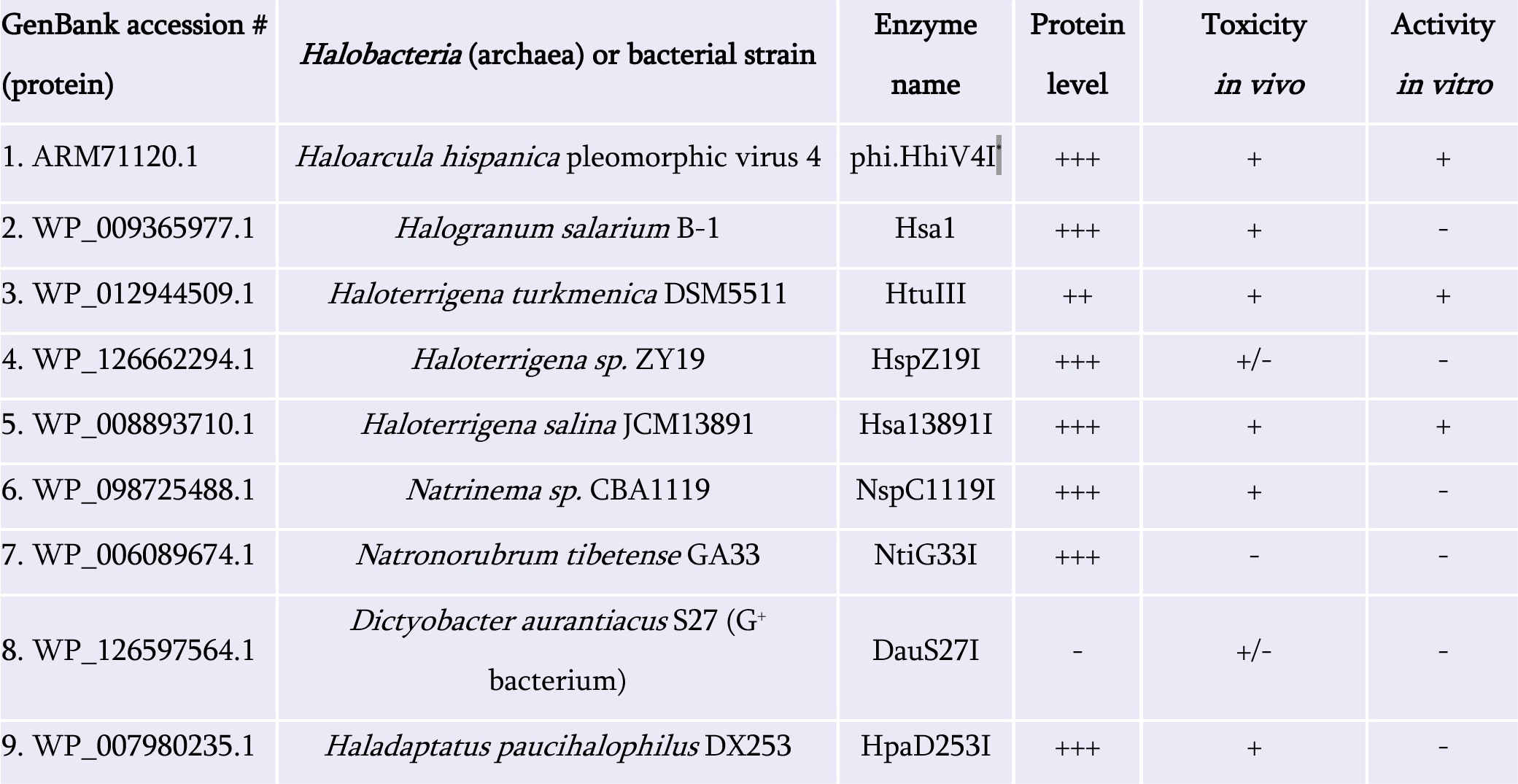
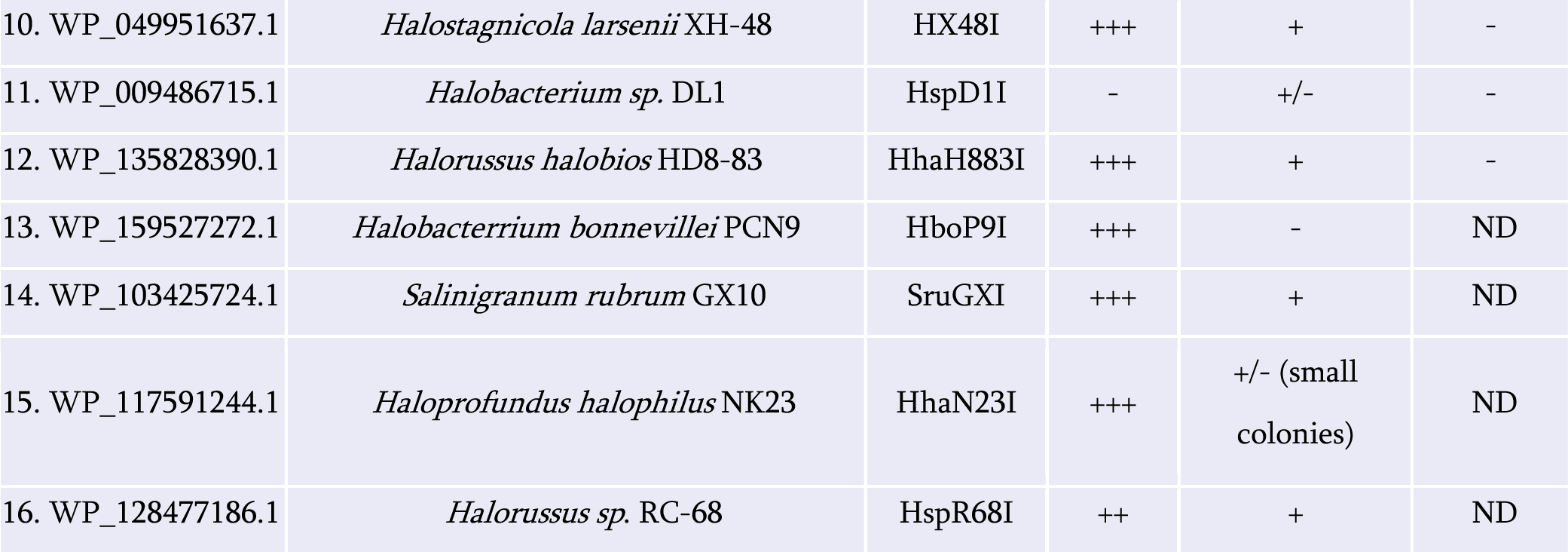
Protein expression, *in vivo* toxicity, and *in vitro* activity of PUA—wH—HNH and wH—HNH endonucleases. The *in vivo* toxic effect of the restriction gene was measured by transformation into Dam^+^ and Dam^-^ *E. coli* competent cells. +, strong restriction (100 to 1000-fold reduction in Dam^+^ cells), +/-, mild restriction (∼10-fold reduction in Dam^+^ cells or formation of sick small colonies), -, no restriction. Protein expression level: +++, 2-10 mg protein per liter of IPTG-induced cells; ++, 1-1.5 mg/L; -, target protein not detected after chitin column purification or in IPTG-induced cell extract. Proteins 1-10: **PUA—wH—HNH** fusions; Proteins 11-16: **wH—HNH** fusions. DauS27I is found in a G^+^ bacterium *Dictyobacter aurantiacus* S27. The other enzymes are found in Archaea (halobacteria). ND, not determined.

Selected enzymes that appeared to be promising as Dam^+^ dependent MDREs, were partially purified, and their activity was tested on Dam^+^ pBR322 or λ DNA. The partially purified HtuIII enzyme (GenBank accession number NC_013743, PUA—wH—HNH fusion) shows a low nicking activity in Mn^2+^ or Co^2+^ buffer **(Suppl. Fig. S8)**.

Digestion of Dam^+^ λ DNA showed some smearing, probably as a result of frequent Dam methylation on the substrate as frequent nickings can collapse dsDNA. DNA run-off sequencing of the partially nicked pBR322 indicated that the nick occurred upstream of the G6mATC site (top strand nicking only (↓NG6mATC-N14-G6mATC). It is not clear whether the partial nicking was due to low enzyme activity. A small amount of linear DNA was also detected in the partial digestion.

Analogous to phi.HhiV4I, HtuIII also preferred Mn^2+^ or Co^2+^ for catalytic activity, suggesting that both enzymes have unique metal ion binding site that is different from typical HNH ββα-metal catalytic domain found in homing endonucleases, Type II REases, Cas9 enzyme, and non-specific endonucleases utilizing Mg^2+^ as cofactor. The preference for Mn^2+^ in cleavage activity is analogous to *E. coli* EcoKMcrA endonuclease and ScoMcrA.

Partially purified Hsa13891I protein (Hsa13891I, NZ_AOIS01000027, 1491 bp, *Haloterrigena salina* JCM 13891 contig_27, whole genome shotgun sequence) (PUA—wH—HNH fusion) also displayed low endonuclease activity on Dam^+^ pBR322 (data not shown). Other fusions did not display apparent *in vitro* cleavage or nicking activity (Table 1).

### wH—GIY-YIG endonucleases

Two wH—GIY-YIG fusion proteins, Ahi29725I (WP_035368356) and Apa233I (WP_026653965) were selected for further study. The proteins occur naturally in *Acholeplasma hippikon* (ATCC29725 strain) and *Acholeplasma palma* (J233 strain), respectively. *Acholeplasma* are bacteria without cell walls in the Mollicutes class with small genomes (1.5 – 1.65 mbp). *Acholeplasma* species are found in animals, insects, and some plants in the environment. Some *Acholeplasma* species are pathogenic and contaminants of mammalian cell cultures. We expressed both proteins in Dam^-^ *E. coli*, and purified the proteins by chromatography through chitin, DEAE, and Heparin columns.

The divalent cation requirement for the Ahi29725I GIY-YIG endonuclease activity was assessed in a medium salt buffer **(Suppl. Fig. S9)**. Ahi29725I is active in Mg^2+^ (1-10 mM) and Mn^2+^ (0.1-10 mM) buffers, and partially active in Co^2+^ (1-10 mM) and Ni^2+^ (1-10 mM) buffers in digestion of pBR322 (Dam^+^). It has a nicking activity in Ca^2+^ (10 mM) buffer. To assess the 6mA-dependent restriction activity, Ahi29725I was also assayed on Dam^-^ pBR322 in the presence of Mg^2+^, Mn^2+^, or Co^2+^in restriction digests. The results showed that the enzyme digested Dam^-^ DNA to a smearing pattern in Mn^2+^ buffer without discrete bands, probably as a result of loss of specificity (data not shown). The exact nature of cut sites in Mn^2+^ buffer remains to be analyzed. It is known that Type II Reases and homing endonucleases (HEases) with GIY-YIG endonuclease domain, preferentially use Mg^2+^ divalent cation as a cofactor.

The purified Apa233I showed similar divalent cation preference as Ahi29725I. The protein is active in restriction buffers with Mg^2+^ or Mn^2+^ (data not shown). Divalent cations Zn^2+^ and Ni^2+^ also support catalytic activity (0.1 mM Zn^2+^ with highest activity). The enzyme displays a low activity in Ca^2+^ buffer. The restriction activity is completely inhibited in EDTA. This is one of the few examples of restriction enzymes that display activity in Zn^2+^ and Ni^2+^ buffers (HpyAV was the first reported Rease that prefers the Ni^2+^ cation (47). Typically the Zn^2+^ cation is required for protein structure folding of HNH restriction endonucleases, even though HNH endonucleases in general accept a broader spectrum of divalent metal cations as co-factors (48-50).

The purified Ahi29725I enzyme was assayed on Dam^+^ and Dam^-^ λ DNA to test modification dependence **(Suppl. Fig. S10)**. Dam^-^ λ DNA was also methylated by Dam methylase (M.Dam) or EcoGII frequent adenine methylase (M.EcoGII) in test tube and used for Ahi29725I digestions.

The Ahi29725I endonuclease generated a partial digestion pattern on Dam^+^ λ DNA and it lacks any cleavage activity on Dam^-^ λ DNA, indicating restriction activity dependent on Dam modification. When Dam^-^ λ DNA was methylated *in vitro* by Dam methylase or M.EcoGII, the modified substrates now became cleavable by Ahi29725I (**Suppl. Fig. S10B**). In control digestion, MboI, DpnII, and Sau3AI are able to cleave Dam^-^ λ DNA, but DpnI cannot. Similarly, Ahi29725I and Apa233I endonucleases are also active on Dam^+^ pBR322 and inactive on Dam^-^ pBR322 (**Fig. 6**). However, high enzyme concentration of Apa233I resulted in non-specific digestion (smearing) on Dam^-^ DNA, which was attributed to the non-specific activity on unmodified DNA.

**Fig. 6.**
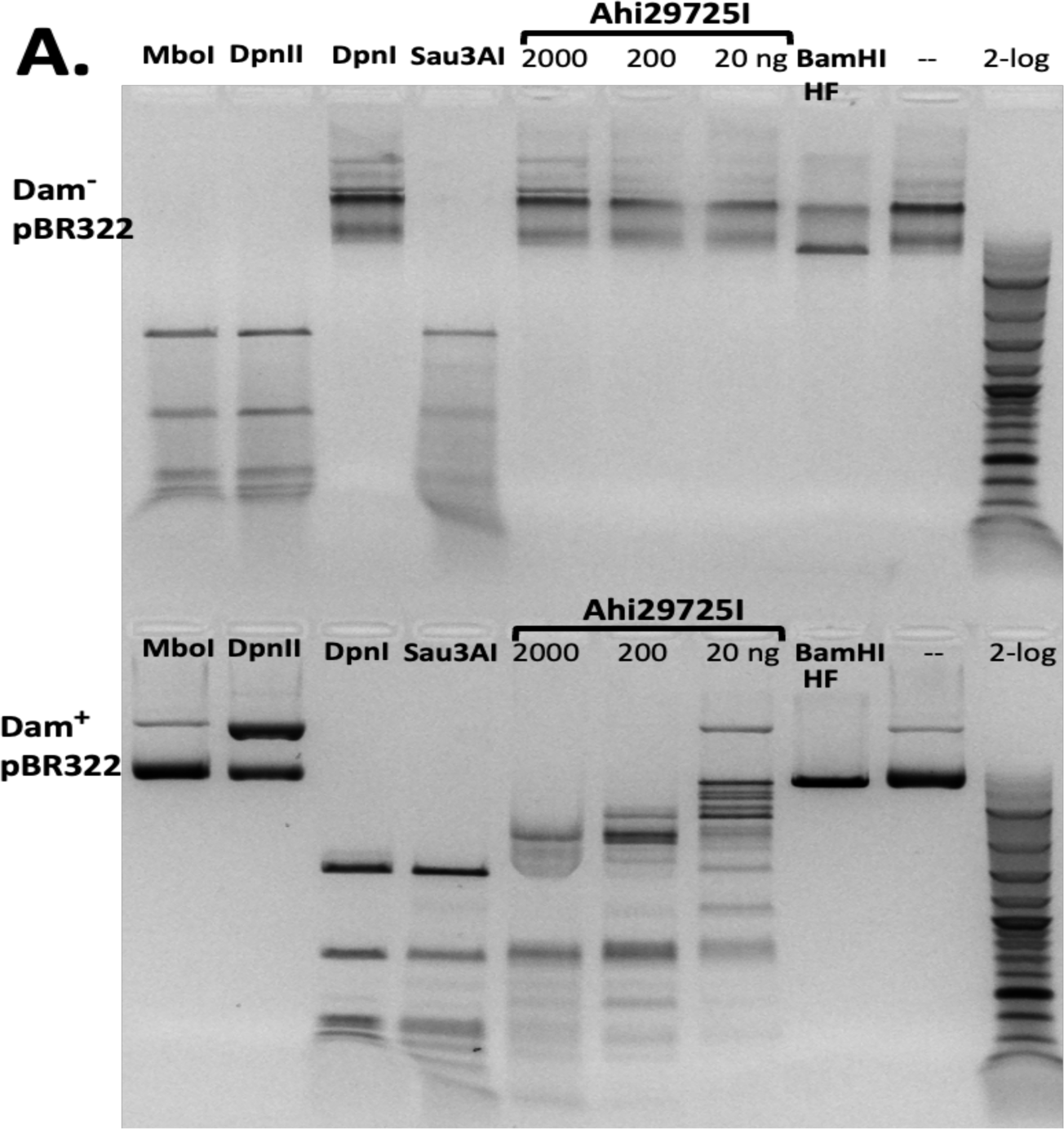
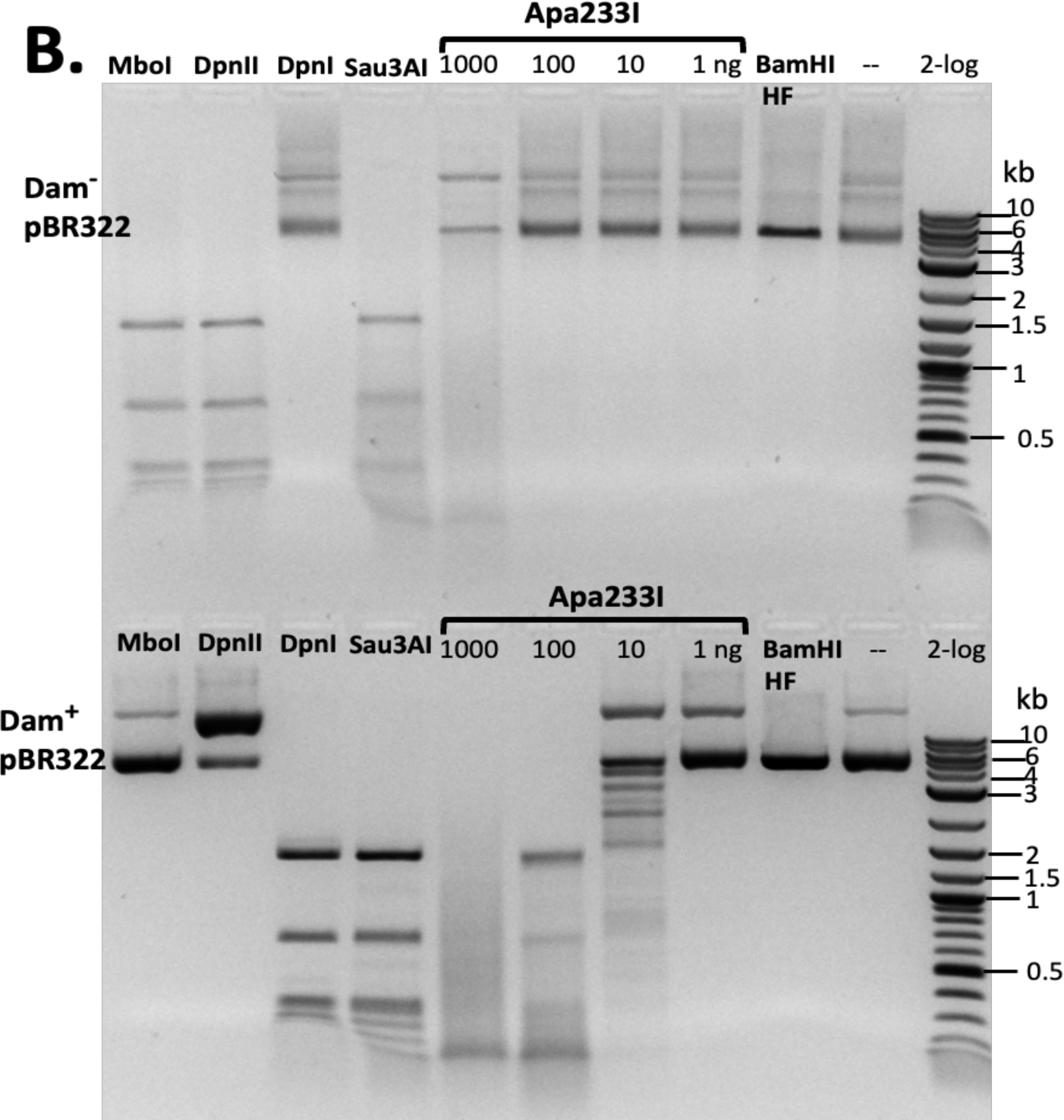
*In vitro* activity of Ahi29725I (A) and Apa233I (B) on Dam^-^ (top) and Dam^+^ (bottom) DNA. Digestion of plasmid DNA (0.5 μg) was done in NEB B2.1 at 37 °C for 1 h. In control digests, MboI and DpnII cleave unmodified GATC sites only; DpnI, a negative control, is unable to cut Dam^-^ DNA. Sau3AI cleaves GATC sites regardless of 6mA methylation. BamHI-HF (GGATCC) was used as an additional control. Apa233I showed non-specific endo/exonuclease activity at high enzyme concentration (at 1 μg). A smeared pattern was detected in both Dam^+^ and Dam^-^ pBR322.

The Ahi29725I and Apa233I digested pBR322 (Dam^+^) DNA was subjected to run-off sequencing with primers annealing near the G6mATC sites. The cleavage distance is variable for both enzymes since all cleavages occur outside of the recognition sequence (i.e. Ahi29725I cleavage outside of G6mATC at N_1-23_) **(Suppl. Fig. S11)**. The cleavage sites of Apa233I were also mapped to the outside of G6mATC sites (**Suppl. Fig. S12**). There is limited sequence specificity at the cleavage positions by the two GIY-YIG endonucleases: most likely cleavage at NN/RN or NN/GN (**Suppl. Fig. S13**).

To test whether Ahi29725I catalyzed DNA cleavage could be directed by adenine methylation outside the G6mATC sequence context, we digested M.EcoGII modified pBR322 DNA (Dam^-^) to see any enhancement of activity due to frequent adenine methylation. M.EcoGII is capable of methylating all adenines in DNA substrate except in polyA tracks (51). Ahi29725I activity was enhanced on M.EcoGII methylated DNA substrate **(Suppl. Fig. S14)**. This result is somewhat surprising in that frequent adenine methylation could enhance Ahi29725I activity.

Three large fragments of Dam^+^ DNA were further digested into smaller fragments after M.EcoGII methylation. However, it was not clear whether the enhanced activity was due to 6mA-dependent relaxed sequence recognition (for example, cleavage near C6mATC star site or S6mATS sites, S = G and C). The enhanced activity on M.EcoGII-modified DNA remains to be characterized in the future.

It is possible that modified star sites C6mATC (G6mATG) could support enzyme binding but cannot activate cleavage (i.e. one enzyme monomer binding to a star site and another monomer binding to a cognate site nearby). Enzyme binding to C6mATC (G6mATG) and G6mATC sites could enhance cleavage activity by dimerization or oligomerization. No significant enhancement of Apa233I activity was observed after 10-100 fold enzyme dilution in digestion of M.EcoGII modified DNA compared to Dam^+^ substrate (data not shown).

In digestion of methylated duplex oligos with a single G6mATC site, it was noted that Ahi29725I preferentially cleaved fully methylated oligos (M+/M+) over hemi-modified substrates (M+/M- or M-/M+). But Apa233I endonuclease was able to cut both fully modified and hemi-modified oligos (**Fig. 4**). This discrepancy of methylation dependence between the two enzymes is still unclear.

### PLD—wH endonucleases

We identified 27 predicted PLD—wH fusion endonucleases in bacterial genomes. Two putative restriction genes from *Anaerolineaceae bacterium* (Aba4572I) and *Chloroflexi bacterium* (CbaI) were cloned into the pTXB1 expression vector. However, upon IPTG induction, no over-expressed proteins were detected. In the gene neighborhood analysis, the Aba45721 ORF resides in a genomic region of 1) DNA MTase (predicted specificity C<u>GATC</u>G, amino-MTase), 2) PLD-wH endonuclease, 3) and 4) hypothetical proteins. If the C<u>GATC</u>G site is methylated to become CG6mATCG which would be a substrate for the PLD—wH endonuclease, it could potentially result in self-restriction. The CbaI enzyme is located in a region with 1) Leu-tRNA ligase, 2) restriction endonuclease, 3) PLD—wH endonuclease, 4) hypothetical protein, 5) dimethyl-menaquinone MTase. Since transformation of Aba4572I and CbaI was less toxic in Dam^-^ cells in non-induced condition **(Fig. 3)**, the lack of expression in Dam^-^ cells is surprising and requires further investigation. Expression of two more PLD-wH fusion proteins containing the conserved catalytic residues HxDxK and HxExK in the PLD endonuclease domain in *E. coli* Dam^-^ cells was not successful, but they showed toxicity in Dam^+^ T7 expression strain (data not shown). More work is necessary to find out the reasons for poor expression.

### Phylogenetic analysis of wH domains of bacterial and Archaea origins

The wH domain sequences from PUA—wH—HNH and wH—HNH endonucleases form a closely related cluster (light blue and deep blue). Similarly, the wH domains from the NTD— wH—NTPase group are closely related to each other and form a cluster (green) (**Suppl. Fig. S15**). The wH domain sequences are phylogenetically related among the PD-(D/E)XK—wH, wH—GIY-YIG, and PLD—wH fusion endonucleases, suggesting they may share a common ancestor in evolution: a similar wH fold was utilized multiple times in combining with endonuclease domains to form the MDREs.. This result is in line with the CLANS analysis of five groups of wH domain sequences of bacteria and Archaea shown in **Fig. 2**.

## Discussion

### wH domain as a sensor of fully methylated ApT in a dsDNA context

The wH domain was first associated with adenine methylation because of its presence in the C-terminal region of *E. coli* and phage T4 Dam methyltransferases (43), and its presence in the adenine methylation dependent REase DpnI. Subsequent work on DpnI showed that the wH domain binds symmetrically adenine methylated ApT DNA sequence, without base flipping. The two methyl groups, which are in close proximity, are bound in a single pocket of the wH domain of DpnI (28). The work on DpnI also showed that the specificity of the domain for the flanking sequence was somewhat relaxed with respect to the Dam G6mATC consensus, to an S6mATS consensus (where S stands for G or C) (29). In this work, we show that the properties as an adenine methylation reader carry over to the fusions with HNH, GIY-YIG and likely also PLD endonuclease domains. If methylation is seen in the ApT context, there is a preference for full-methylation except Apa233I (**Fig. 3 and Fig. 4**). Our work shows that in all tested fusion proteins, the wH domain can operate as an adenine methylation reader for the G6mATC context. Work on the wH—GIY-YIG endonucleases further indicates that additional cleavage sites are created when Dam methylated DNA is hypermethylated by M.EcoII (**Suppl. Fig. S14**). Hence, the wH domains of the wH—GIY-YIG endonucleases also suggest that the G6mATC preference may be relaxed, albeit, according to the digestion experiments with synthetic DNA, not necessarily in the same way.

The identification of prokaryotic winged helix domains as sensors of adenine methylation contrasts with the role of some eukaryotic winged helix domains as sensors of non-methylated CpG (39-41). Superposition of the winged helix domains of prokaryotic DpnI (28) and eukaryotic KAT6A (a.k.a. Histone Acetyltransferase KAT6A, Lysine Acetyltransferase 6A, Zinc Finger Protein 220, MYST-3) (41) shows that the dsDNA molecules are bound to opposite faces of the wH domain **(Suppl. Fig. S16)**, indicating that the two DNA binding modes have likely evolved independently, for needs that are characteristic for prokaryotes (sensing of Dam methylation), and eukaryotes (sensing of the absence of CpG methylation).

### Cooperation with endonuclease domains

For most NTP-independent MDREs, there is a clear division of labor between the modification reader and endonuclease domains. The former recruits the enzyme to modification sites, and the latter cleaves the DNA at a distance from the recognition site, that is likely defined by the length of the linker that connects the two domains. The nuclease domain has generally low or only a very relaxed sequence specificity, and it is not modification specific. How the modification sensor domain keeps the activity of the nuclease domain in check is not well understood. However, in some cases it can be shown that the linker has an inhibitory role for the endonuclease that is only relieved once modified DNA is bound to the reader domain and the complex reorganizes structurally (19). The PUA—wH—HNH (phi.HhiV4I) and wH— GIY-YIG (Ahi29725I and Apa233I) that were tested by run-off sequencing are consistent with this expectation. As for the cytosine modification specific MDREs, cleavage occurred always at a distance from the recognition sequence except the BisI family REases (e.g. Eco15I and NhoI) that cut within the recognition sequence G5mCNGC.

Among the wH fusion endonucleases, DpnI is the exception to the rule that cleavage occurs always outside of, and not within the recognition sequence. Mechanically, DpnI DNA cleavage within the recognition sequence is a consequence of the fact that the endonuclease domain has separate sequence and modification specificity (29). In this scenario, the role of the wH domain is similar to the role of the extra specificity domain in Type IIE restriction endonucleases, except that the target sequence contains a modified base. Run-off sequencing shows that FcyTI and Psp4BI cleave inside the recognition sequence, like DpnI, pointing to separate sequence and modification specificity of the catalytic PD-(D/E)XK domain. Given the high sequence conservation of the PD-(D/E)XK—wH endonuclease family, it is likely that this property is general for the entire family.

### Sensors for N6-methyladenine in DNA

Despite the many roles of DNA adenine methylation in prokaryotes (and eukaryotic organelles), the repertoire of reader domains for 6mA in DNA is still surprisingly limited (23). Perhaps the best-known adenine methylation sensors are YTH (52) domains, which belong to (or are related to) the PUA superfamily domains. The PUA superfamily domains are believed to flip the modified 2′-deoxynucleotide in DNA (20), or to bind a single nucleotide in RNA in the reader pocket (53,54). Therefore, at least when acting in isolation, they can be considered as sensors of a single modified adenine. Consistent with this role, most YTH domains sense adenine methylation in RNA, rather than DNA. However, some YTH and ASCH domains in prokaryotes are considered as DNA adenine methylation sensors (23). Our preliminary data on expression of Yth-PD-(D/E)xK fusion endonuclease suggests it is toxic to Dam^+^ *E. coli* host (SYX, unpublished). For the ASCH domains, this remains to be experimentally shown, since currently only a 4mC reader role is experimentally supported (24). Apart from YTH and ASCH domains, HARE-HTH and RAMA domains have also been suggested to serve as readers of adenine methylation in DNA (43). The HARE-HTH domains are related to winged helix domains, but have an extra helix inserted into the HTH motif of the winged helix domain, which in the light of this manuscript would be consistent with a role as adenine methylation sensors. However, recent analysis suggests that they are more likely to sense cytosine modifications (55). Unlike the HARE-HTH domains, the RAMA (<u>R</u>estriction enzyme <u>A</u>denine Methylation <u>A</u>ssociated) domains are unrelated to the wH domain in fold (56). For the RAMA domain containing MPND protein, there is some biochemical evidence for adenine methylation sensing (57). However, a preference for adenine methylated DNA could not be experimentally confirmed (56). We noticed the occurrence of RAMA—Mrr catalytic domain—NTPase and GIY-YIG—RAMA fusions in prokaryotes, which might indicate that the RAMA domain is used similarly to the wH domain in these fusions. Given the scarcity of readers of 6mA in DNA, our demonstration that the wH domain is used widely as a sensor of adenine methylation in both strands in ApT context considerably extends the landscape of reader domains for 6mA in DNA.

## Acknowledgement

Part of this work was supported by a grant from the Polish government project NAWA (PPI/APM/2018/1/00034/U/001). This work was also supported by New England Biolabs, Inc (NEB). We thank Andy Gardner, Tom Evans, Rich Roberts, Jim Ellard, and Salvatore Russello for continued support. We thank Drs. Honorata Czapinska and Lise Raleigh for critical reading of the manuscript. The publication cost (open access fee) is paid for by NEB.

## References

1. Miyazono, K., Furuta, Y., Watanabe-Matsui, M., Miyakawa, T., Ito, T., Kobayashi, I. and Tanokura, M. (2014) A sequence-specific DNA glycosylase mediates restriction-modification in Pyrococcus abyssi. Nat Commun, 5, 3178.

2. Pingoud, A., Fuxreiter, M., Pingoud, V. and Wende, W. (2005) Type II restriction endonucleases: structure and mechanism. Cell Mol Life Sci, 62, 685–707.

3. Kosinski, J., Feder, M. and Bujnicki, J.M. (2005) The PD-(D/E)XK superfamily revisited: identification of new members among proteins involved in DNA metabolism and functional predictions for domains of (hitherto) unknown function. BMC Bioinformatics, 6, 172.

4. Bujnicki, J.M. and Rychlewski, L. (2001) Grouping together highly diverged PD-(D/E)XK nucleases and identification of novel superfamily members using structure-guided alignment of sequence profiles. J Mol Microbiol Biotechnol, 3, 69–72.

5. Jablonska, J., Matelska, D., Steczkiewicz, K. and Ginalski, K. (2017) Systematic classification of the His-Me finger superfamily. Nucleic Acids Res, 45, 11479–11494.

6. Wu, C.C., Lin, J.L.J. and Yuan, H.S. (2020) Structures, Mechanisms, and Functions of His-Me Finger Nucleases. Trends Biochem Sci, 45, 935–946.

7. Pommer, A.J., Cal, S., Keeble, A.H., Walker, D., Evans, S.J., Kuhlmann, U.C., Cooper, A., Connolly, B.A., Hemmings, A.M., Moore, G.R. et al. (2001) Mechanism and cleavage specificity of the H-N-H endonuclease colicin E9. J Mol Biol, 314, 735–749.

8. Flick, K.E., Jurica, M.S., Monnat, R.J., Jr. and Stoddard, B.L. (1998) DNA binding and cleavage by the nuclear intron-encoded homing endonuclease I-PpoI. Nature, 394, 96–101.

9. Sokolowska, M., Czapinska, H. and Bochtler, M. (2009) Crystal structure of the beta beta alpha-Me type II restriction endonuclease Hpy99I with target DNA. Nucleic Acids Res, 37, 3799–3810.

10. Sokolowska, M., Czapinska, H. and Bochtler, M. (2011) Hpy188I-DNA pre- and post-cleavage complexes--snapshots of the GIY-YIG nuclease mediated catalysis. Nucleic Acids Res, 39, 1554–1564.

11. Grazulis, S., Manakova, E., Roessle, M., Bochtler, M., Tamulaitiene, G., Huber, R. and Siksnys, V. (2005) Structure of the metal-independent restriction enzyme BfiI reveals fusion of a specific DNA-binding domain with a nonspecific nuclease. Proc Natl Acad Sci U S A, 102, 15797–15802.

12. Sasnauskas, G., Zakrys, L., Zaremba, M., Cosstick, R., Gaynor, J.W., Halford, S.E. and Siksnys, V. (2010) A novel mechanism for the scission of double-stranded DNA: BfiI cuts both 3’-5’ and 5’-3’ strands by rotating a single active site. Nucleic Acids Res, 38, 2399–2410.

13. Lutz, T., Flodman, K., Copelas, A., Czapinska, H., Mabuchi, M., Fomenkov, A., He, X., Bochtler, M. and Xu, S.Y. (2019) A protein architecture guided screen for modification dependent restriction endonucleases. Nucleic Acids Res, 47, 9761–9776.

14. Cohen-Karni, D., Xu, D., Apone, L., Fomenkov, A., Sun, Z., Davis, P.J., Kinney, S.R., Yamada-Mabuchi, M., Xu, S.Y., Davis, T. et al. (2011) The MspJI family of modification-dependent restriction endonucleases for epigenetic studies. Proc Natl Acad Sci U S A, 108, 11040–11045.

15. Kazrani, A.A., Kowalska, M., Czapinska, H. and Bochtler, M. (2014) Crystal structure of the 5hmC specific endonuclease PvuRts1I. Nucleic Acids Res, 42, 5929–5936.

16. Shao, C., Wang, C. and Zang, J. (2014) Structural basis for the substrate selectivity of PvuRts1I, a 5-hydroxymethylcytosine DNA restriction endonuclease. Acta Crystallogr D Biol Crystallogr, 70, 2477–2486.

17. Borgaro, J.G. and Zhu, Z. (2013) Characterization of the 5-hydroxymethylcytosine-specific DNA restriction endonucleases. Nucleic Acids Res, 41, 4198–4206.

18. Janosi, L., Yonemitsu, H., Hong, H. and Kaji, A. (1994) Molecular cloning and expression of a novel hydroxymethylcytosine-specific restriction enzyme (PvuRts1I) modulated by glucosylation of DNA. J Mol Biol, 242, 45–61.

19. Pastor, M., Czapinska, H., Helbrecht, I., Krakowska, K., Lutz, T., Xu, S.Y. and Bochtler, M. (2021) Crystal structures of the EVE-HNH endonuclease VcaM4I in the presence and absence of DNA. Nucleic Acids Res, 49, 1708–1723.

20. Hosford, C.J., Bui, A.Q. and Chappie, J.S. (2020) The structure of the Thermococcus gammatolerans McrB N-terminal domain reveals a new mode of substrate recognition and specificity among McrB homologs. J Biol Chem, 295, 743–756.

21. Iyer, L.M., Burroughs, A.M. and Aravind, L. (2006) The ASCH superfamily: novel domains with a fold related to the PUA domain and a potential role in RNA metabolism. Bioinformatics, 22, 257–263.

22. Xu, D., Shao, J., Song, H. and Wang, J. (2020) The YTH Domain Family of N6-Methyladenosine “Readers” in the Diagnosis and Prognosis of Colonic Adenocarcinoma. Biomed Res Int, 2020, 9502560.

23. Iyer, L.M., Zhang, D. and Aravind, L. (2016) Adenine methylation in eukaryotes: Apprehending the complex evolutionary history and functional potential of an epigenetic modification. Bioessays, 38, 27–40.

24. Stanislauskiene, R., Laurynenas, A., Rutkiene, R., Aucynaite, A., Tauraite, D., Meskiene, R., Urbeliene, N., Kaupinis, A., Valius, M., Kaliniene, L. et al. (2020) YqfB protein from Escherichia coli: an atypical amidohydrolase active towards N(4)-acylcytosine derivatives. Sci Rep, 10, 788.

25. Horton, J.R., Wang, H., Mabuchi, M.Y., Zhang, X., Roberts, R.J., Zheng, Y., Wilson, G.G. and Cheng, X. (2014) Modification-dependent restriction endonuclease, MspJI, flips 5-methylcytosine out of the DNA helix. Nucleic Acids Res, 42, 12092-12101.

26. Czapinska, H., Kowalska, M., Zagorskaite, E., Manakova, E., Slyvka, A., Xu, S.Y., Siksnys, V., Sasnauskas, G. and Bochtler, M. (2018) Activity and structure of EcoKMcrA. Nucleic Acids Res, 46, 9829–9841.

27. Slyvka, A., Zagorskaite, E., Czapinska, H., Sasnauskas, G. and Bochtler, M. (2019) Crystal structure of the EcoKMcrA N-terminal domain (NEco): recognition of modified cytosine bases without flipping. Nucleic Acids Res, 47, 11943–11955.

28. Mierzejewska, K., Siwek, W., Czapinska, H., Kaus-Drobek, M., Radlinska, M., Skowronek, K., Bujnicki, J.M., Dadlez, M. and Bochtler, M. (2014) Structural basis of the methylation specificity of R.DpnI. Nucleic Acids Res, 42, 8745–8754.

29. Siwek, W., Czapinska, H., Bochtler, M., Bujnicki, J.M. and Skowronek, K. (2012) Crystal structure and mechanism of action of the N6-methyladenine-dependent type IIM restriction endonuclease R.DpnI. Nucleic Acids Res, 40, 7563–7572.

30. Brennan, R.G. (1993) The winged-helix DNA-binding motif: another helix-turn-helix takeoff. Cell, 74, 773–776.

31. Gajiwala, K.S. and Burley, S.K. (2000) Winged helix proteins. Curr Opin Struct Biol, 10, 110–116.

32. Lai, E., Clark, K.L., Burley, S.K. and Darnell, J.E., Jr. (1993) Hepatocyte nuclear factor 3/fork head or “winged helix” proteins: a family of transcription factors of diverse biologic function. Proc Natl Acad Sci U S A, 90, 10421–10423.

33. Teichmann, M., Dumay-Odelot, H. and Fribourg, S. (2012) Structural and functional aspects of winged-helix domains at the core of transcription initiation complexes. Transcription, 3, 2–7.

34. Schwartz, T., Behlke, J., Lowenhaupt, K., Heinemann, U. and Rich, A. (2001) Structure of the DLM-1-Z-DNA complex reveals a conserved family of Z-DNA-binding proteins. Nat Struct Biol, 8, 761–765.

35. Tang, Q., Rigby, R.E., Young, G.R., Hvidt, A.K., Davis, T., Tan, T.K., Bridgeman, A., Townsend, A.R., Kassiotis, G. and Rehwinkel, J. (2021) Adenosine-to-inosine editing of endogenous Z-form RNA by the deaminase ADAR1 prevents spontaneous MAVS-dependent type I interferon responses. Immunity, 54, 1961–1975 e1965.

36. Wah, D.A., Hirsch, J.A., Dorner, L.F., Schildkraut, I. and Aggarwal, A.K. (1997) Structure of the multimodular endonuclease FokI bound to DNA. Nature, 388, 97–100.

37. Gajiwala, K.S., Chen, H., Cornille, F., Roques, B.P., Reith, W., Mach, B. and Burley, S.K. (2000) Structure of the winged-helix protein hRFX1 reveals a new mode of DNA binding. Nature, 403, 916–921.

38. Wolberger, C. and Campbell, R. (2000) New perch for the winged helix. Nat Struct Biol, 7, 261–262.

39. Becht, D.C., Klein, B.J., Kanai, A., Jang, S.M., Cox, K.L., Zhou, B.R., Phanor, S.K., Zhang, Y., Chen, R.W., Ebmeier, C.C. et al. (2023) MORF and MOZ acetyltransferases target unmethylated CpG islands through the winged helix domain. Nat Commun, 14, 697.

40. Stielow, B., Zhou, Y., Cao, Y., Simon, C., Pogoda, H.M., Jiang, J., Ren, Y., Phanor, S.K., Rohner, I., Nist, A. et al. (2021) The SAM domain-containing protein 1 (SAMD1) acts as a repressive chromatin regulator at unmethylated CpG islands. Sci Adv, 7.

41. Weber, L.M., Jia, Y., Stielow, B., Gisselbrecht, S.S., Cao, Y., Ren, Y., Rohner, I., King, J., Rothman, E., Fischer, S. et al. (2023) The histone acetyltransferase KAT6A is recruited to unmethylated CpG islands via a DNA binding winged helix domain. Nucleic Acids Res, 51, 574–594.

42. Lu, X., Huang, F., Cheng, R. and Zhu, B. (2023) A Unique m6A-Dependent Restriction Endonuclease from an Archaeal Virus. Microbiol Spectr, e0426222.

43. Marinus, M.G. and Casadesus, J. (2009) Roles of DNA adenine methylation in host-pathogen interactions: mismatch repair, transcriptional regulation, and more. FEMS Microbiol Rev, 33, 488–503.

44. Boye, E. and Lobner-Olesen, A. (1990) The role of dam methyltransferase in the control of DNA replication in E. coli. Cell, 62, 981–989.

45. Au, K.G., Welsh, K. and Modrich, P. (1992) Initiation of methyl-directed mismatch repair.J Biol Chem, 267, 12142–12148.

46. Josephs, E.A., Zheng, T. and Marszalek, P.E. (2015) Atomic force microscopy captures the initiation of methyl-directed DNA mismatch repair. DNA Repair (Amst), 35, 71–84.

47. Chan, S.H., Opitz, L., Higgins, L., O’Loane, D. and Xu, S.Y. (2010) Cofactor requirement of HpyAV restriction endonuclease. PLoS One, 5, e9071.

48. Walker, D.C., Georgiou, T., Pommer, A.J., Walker, D., Moore, G.R., Kleanthous, C. and James, R. (2002) Mutagenic scan of the H-N-H motif of colicin E9: implications for the mechanistic enzymology of colicins, homing enzymes and apoptotic endonucleases. Nucleic Acids Res, 30, 3225–3234.

49. Galburt, E.A., Chevalier, B., Tang, W., Jurica, M.S., Flick, K.E., Monnat, R.J., Jr. and Stoddard, B.L. (1999) A novel endonuclease mechanism directly visualized for I-PpoI. Nat Struct Biol, 6, 1096–1099.

50. Meiss, G., Franke, I., Gimadutdinow, O., Urbanke, C. and Pingoud, A. (1998) Biochemical characterization of Anabaena sp. strain PCC 7120 non-specific nuclease NucA and its inhibitor NuiA. Eur J Biochem, 251, 924–934.

51. Murray, I.A., Morgan, R.D., Luyten, Y., Fomenkov, A., Correa, I.R., Jr., Dai, N., Allaw, M.B., Zhang, X., Cheng, X. and Roberts, R.J. (2018) The non-specific adenine DNA methyltransferase M.EcoGII. Nucleic Acids Res, 46, 840–848.

52. Liao, S., Sun, H. and Xu, C. (2018) YTH Domain: A Family of N(6)-methyladenosine (m(6)A) Readers. Genomics Proteomics Bioinformatics, 16, 99–107.

53. Li, F., Zhao, D., Wu, J. and Shi, Y. (2014) Structure of the YTH domain of human YTHDF2 in complex with an m(6)A mononucleotide reveals an aromatic cage for m(6)A recognition. Cell Res, 24, 1490–1492.

54. Xu, C., Wang, X., Liu, K., Roundtree, I.A., Tempel, W., Li, Y., Lu, Z., He, C. and Min, J. (2014) Structural basis for selective binding of m6A RNA by the YTHDC1 YTH domain. Nat Chem Biol, 10, 927–929.

55. Aravind, L. and Iyer, L.M. (2012) The HARE-HTH and associated domains: novel modules in the coordination of epigenetic DNA and protein modifications. Cell Cycle, 11, 119–131.

56. Yang, M., Li, X., Tian, Z., Ma, L., Ma, J., Liu, Y., Shang, G., Liang, A., Wu, W. and Chen, Z. (2023) Structures of MPND Reveal the Molecular Recognition of Nucleosomes. Int J Mol Sci, 24.

57. Kweon, S.M., Chen, Y., Moon, E., Kvederaviciute, K., Klimasauskas, S. and Feldman, D.E. (2019) An Adversarial DNA N(6)-Methyladenine-Sensor Network Preserves Polycomb Silencing. Mol Cell, 74, 1138–1147 e1136.

